# Diversity of the non-biting midges (Diptera: Chironomidae) in continental salt marshes in Serbia

**DOI:** 10.1101/2024.04.10.588933

**Authors:** Olivera Stamenković, Dubravka Čerba, Aca Đurđević, Miran Koh

## Abstract

Continental salt marshes represent specific inland saltwater bodies with unique ecological characteristics a fact that is mirrored by specific flora and fauna. However, there is still limited data on their biota in Serbia, which is especially true for aquatic macroinvertebrates, including chironomids (Diptera: Chironomidae). Here we investigated diversity and seasonal variations of chironomid community in six salt marshes distributed in the northern and southern part of Serbia. We recorded a total of 25 species, of which two are new for the Serbian chironomid fauna. Most of the recorded species are common in freshwaters and several of them are halotolerant. Chironomid community structure fluctuated in relation to the seasons. The highest diversity was recorded in the spring and summer months for the majority of studied salt marshes. The present findings should contribute to the knowledge of the faunistic composition of salt marshes in Serbia and provide a basis for future distributional and ecological studies of chironomids.

## Introduction

Continental salt marshes and soda pans represent specific inland saltwater bodies with unique ecological characteristics, which makes them the most vulnerable aquatic habitats in Europe (Boros et al., 2014). In Europe, the Carpathian basin is particularly rich in these types of habitats, with natural salt marshes found in Austria, Hungary, and Serbia. At the EU level, these habitats are classified as “Pannonian salt steppes and salt marshes” habitat type (Vidaković et al., 2019). They are listed in Annex I of the EU Habitat Directive (93/43/EEC) and are part of the Natura 2000 Network. In Serbia, the Pannonian salt marshes are located in the northern part of the country, Vojvodina Province. Besides these types of salt marshes, a few isolated continental salt marshes are located in southern Serbia, in the valleys of the Južna Morava and Toplica rivers (Zlatković et al., 2014).

Continental salt marshes encompass various types of lentic water bodies ranging from small stagnant waters to large lakes, but the common feature of all types of these saline water bodies is their shallow depth. Besides the shallowness, some of the common features of the continental salt marshes are high conductivity with mainly Na^+^–HCO_3_^-^ ionic dominance, high water pH value (9-10), high seasonal water-level fluctuations, and high daily and seasonal temperature variations (Lengyel et al., 2016). Some other environmental parameters, such as salinity, can also show strong annual variations, changing over hypo-, meso- and hypersaline limits (Boros et al., 2014). Thus, continental salt marshes are inhabited by unique flora and fauna that can tolerate extreme environmental conditions (Gavrilović et al., 2018). Salt marshes have not been studied extensively in Serbia, and therefore, there is still limited data on their biota. Previous studies have majorly focused on the diversity of diatoms and crustaceans in Pannonian salt marshes in Vojvodina Province (Lukić et al., 2012; Gavrilović et al., 2018; Vidaković et al., 2019), while saltmarshes in the southern part of Serbia have been extensively studied only in terms of their floristic composition (Zlatković et al., 2014), with scarce published data on the faunistic composition of some of the localities, such as data on avifauna in Lalinačka salt marsh (Nikolić & Ilić, 2021). Macroinvertebrate communities of the continental salt marshes in Europe have been generally poorly studied (Boda et al., 2019). In Serbia, except some information on aquatic species of Coleoptera in Pannonian salt marshes in the Vojvodina Province (Gavrilović et al., 2018), there is no published data on any other groups of aquatic macroinvertebrates in salt marshes.

Non-biting midges (Diptera: Chironomidae) are a widely distributed insect family which represents the most diverse and often the most abundant group of macroinvertebrates in both lotic and lentic ecosystems (Ferrington, 2008). This group also plays an important role in ecosystem functioning due to their intermediate position in the food-web and their key role in detritus processing in freshwaters (da Silva & Henry, 2018). The absence of fish in continental salt marshes allows development of diverse chironomid assemblages that represent a mixture of halobiont and halophil species, but also eurytopic species with high geographical distribution can be abundant (Boda et al., 2019). Previous research on chironomid communities on the Balkan Peninsula has been conducted majorly in different freshwater ecosystems (Milošević et al., 2011; Płóciennik & Pešić, 2012; Bitušík & Trnkova, 2019; Popović et al., 2016, 2022; Dorić et al., 2020; Gadawski et al., 2022; Čerba et al., 2023; Ergović et al., 2023) with some records on chironomid species in coastal regions in Montenegro and Croatia (Płóciennik et al., 2012; Čerba et al., 2020). To date, chironomid fauna in continental salt marshes on Balkan Peninsula has not been investigated. Moreover, chironomid fauna in Serbia is still not explored sufficiently. To date only three extensive faunistic studies on chironomid communities in Serbia have been published (Janković, 1978, 1985; Milošević et al., 2011). Besides these faunistic studies, the data on chironomid fauna comes from several ecological studies on chironomids (Milošević et al., 2012, 2013, 2018; Popović et al., 2016, 2022).

The aim of the present study is to contribute to the knowledge on diversity and distribution of chironomid larvae in Serbia while contributing at the same time to the knowledge on animal diversity of continental salt marshes.

## Material and methods

### Study sites

The study encompasses a total of six salt marshes (study sites), of which three are Pannonian salt marshes in the northern part of Serbia declared as protected areas: Special nature reserve “Slano Kopovo” (SK), Special nature reserve “Okanj bara” (OK) and Nature Park “Rusanda” (RU; Fig. 1). These three salt marshes аrе paleo-meanders of the Tisa River situated on its left bank about 15 km northwest of the city of Zrenjanin (Fig. 1). The maximum depth of these saline lakes does not exceed 1.5 m, while the sizes of surface area are around 1.5 km^2^ (Fig. 2d, e and f). The remaining three study sites are small salt marshes located in the southeastern part of the country, in the valleys of the Južna Morava and Toplica Rivers (Fig. 1). Two of them belong to Lalinac salt marsh area in the vicinity of the village of Lalinac near the city of Niš (Lalinačka salt marsh, LA and Oblačinska salt marsh, OB; Fig. 1), while the remaining one belongs to Bresničić salt marsh area in the vicinity of the village of Bresničić near the city of Prokuplje (Bresničićka salt marsh, BR; Fig. 1). These salt marshes are small water bodies whose surface areas do not exceed 0.02 km^2^ (Fig. 2a, b and c). Two of them are declared as protected areas: Natural monument “Lalinačka slatina” and Natural monument “Bresničićka slatina”.

**Fig. 1.**
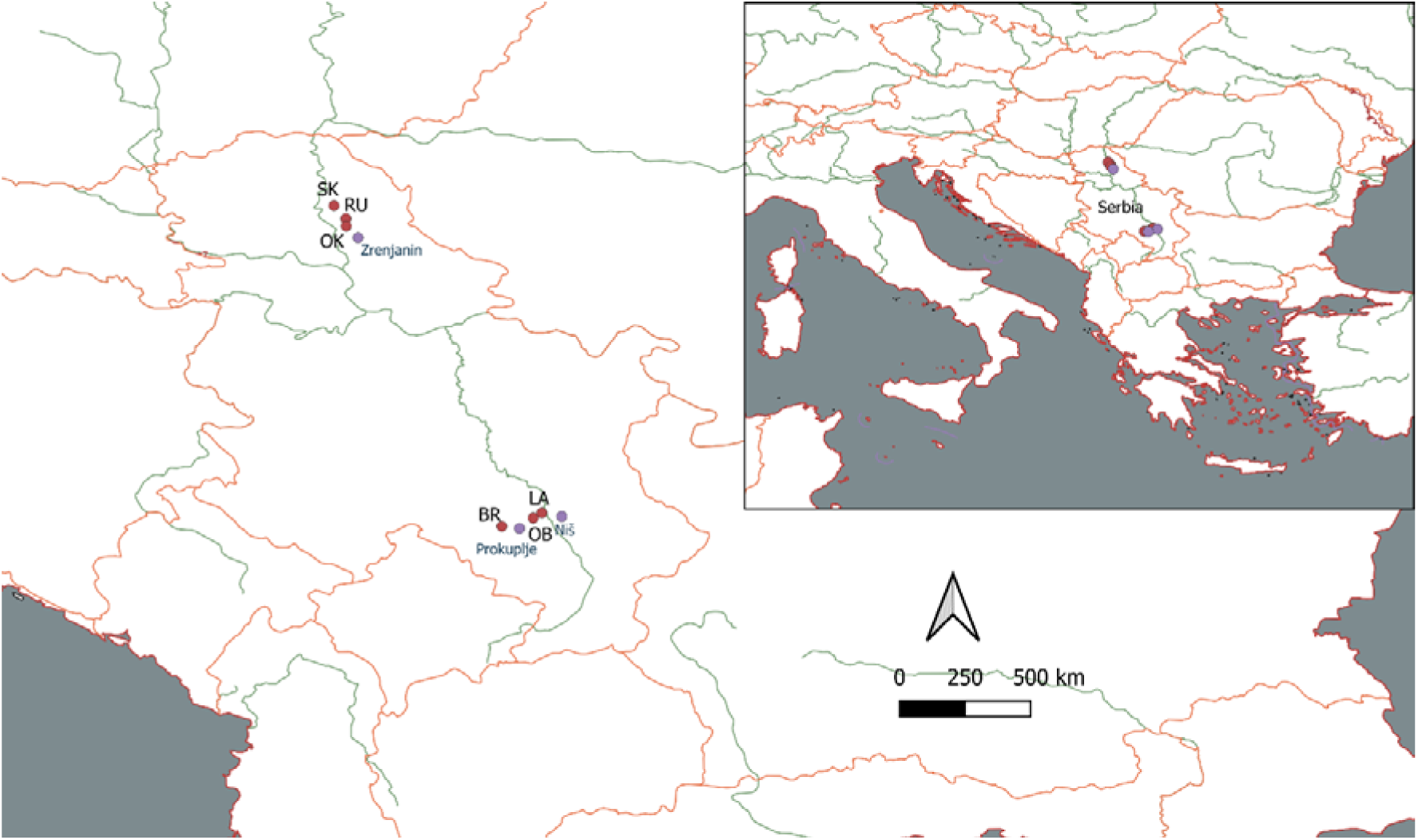
Map of the study sites: Lalinačka salt marsh, LA; Oblačinska salt marsh, OB; Bresničićka salt marsh, BR; Okanj bara, OK; Slano Kopovo, SK; Rusanda, RU.

**Fig. 2.**
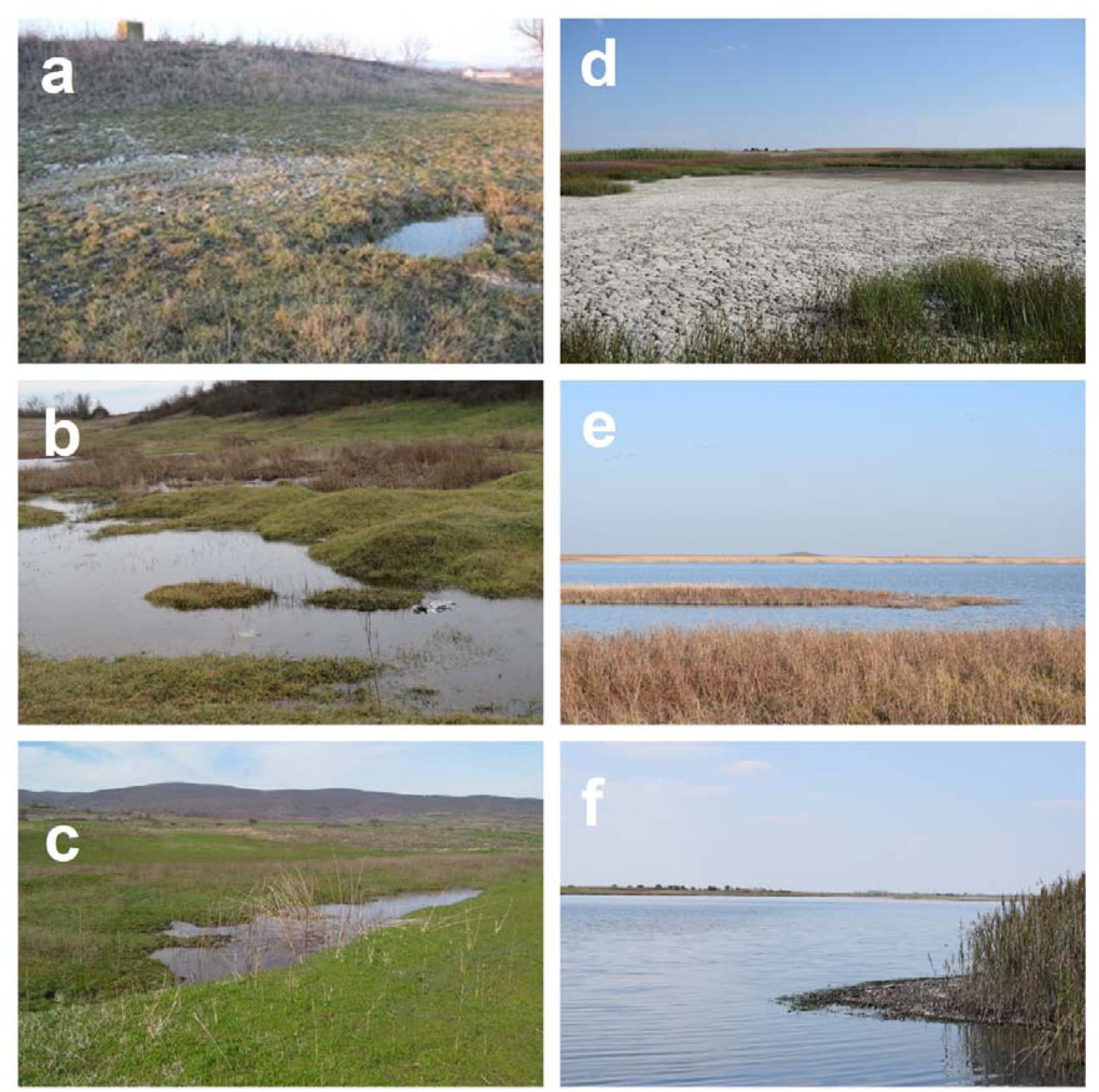
Photographs of the study sites: Oblačinska salt marsh (a), Bresničićka salt marsh (b), Lalinačka salt marsh (c), Okanj bara (d), Slano Kopovo (e), Rusanda (f)

### Sampling and analysis

Sampling campaign was performed from December 2016 to November 2017. Samples were collected at each salt marsh (hereinafter study site) every month except at Lalinačka salt marsh, which was not sampled during summer as it had dried up in late spring. Okanj bara, Slano Kopovo and Oblačinska salt marsh were sampled only once during summer before they completely dried up.

Given that not all study sites were sampled every month, the results below are presented for seasons. Benthic samples were collected by using a hand net of 250 µm mesh size. A total of three subsamples were taken from the most common substrates at each study site, except at Oblačinska salt marsh where only one sample was taken due to the small size of this study site (Fig. 2a). The collected material was sorted out and preserved in 70% ethanol. Chironomid larvae were identified to the species or genus level using relevant taxonomic keys (Moller Pillot, 1984a, b, 2009; Schmid, 1993; Andersen et al., 2013; Rossaro & Lencioni, 2015). The abundance of chironomids was expressed as the number of individuals per m^2^. Chironomid diversity was expressed as species richness, S (number of taxa in a sample) and by calculating Shannon diversity, H (Shannon and Weaver, 1963). To visualize the prevalence of the chironomid taxa across study sites during each season we created pie charts using “ggplot2” package (Wickham, 2016) in R version 4.3.1 (R Core Team, 2023).

## Results

### Species richness and diversity of chironomid communities

Overall, a total of 25 chironomid taxa were recorded across all study sites and throughout all four seasons (Table 1). The richest site was Bresničićka salt marsh where 17 taxa were recorded, followed by Oblačinska salt marsh with 11 taxa, and Lalinačka salt marsh with 9 taxa (Table 1). Lower species richness was recorded at study sites in the northern part of the country: 5 taxa at Slano Kopovo, 4 at Okanj bara, and 3 at Rusanda. Shannon diversity was the highest at Lalinačka salt marsh and Slano Kopovo since certain taxa dominated in the abundance at the remaining localities. According to the published data for chironomids in Serbia, among 25 chironomid taxa recorded in our study two are new for the chironomid fauna of Serbia: *Microchironomus* cf. *deribae* and *Tanytarsus* gr. *pallidicornis*.

**Table 1.**
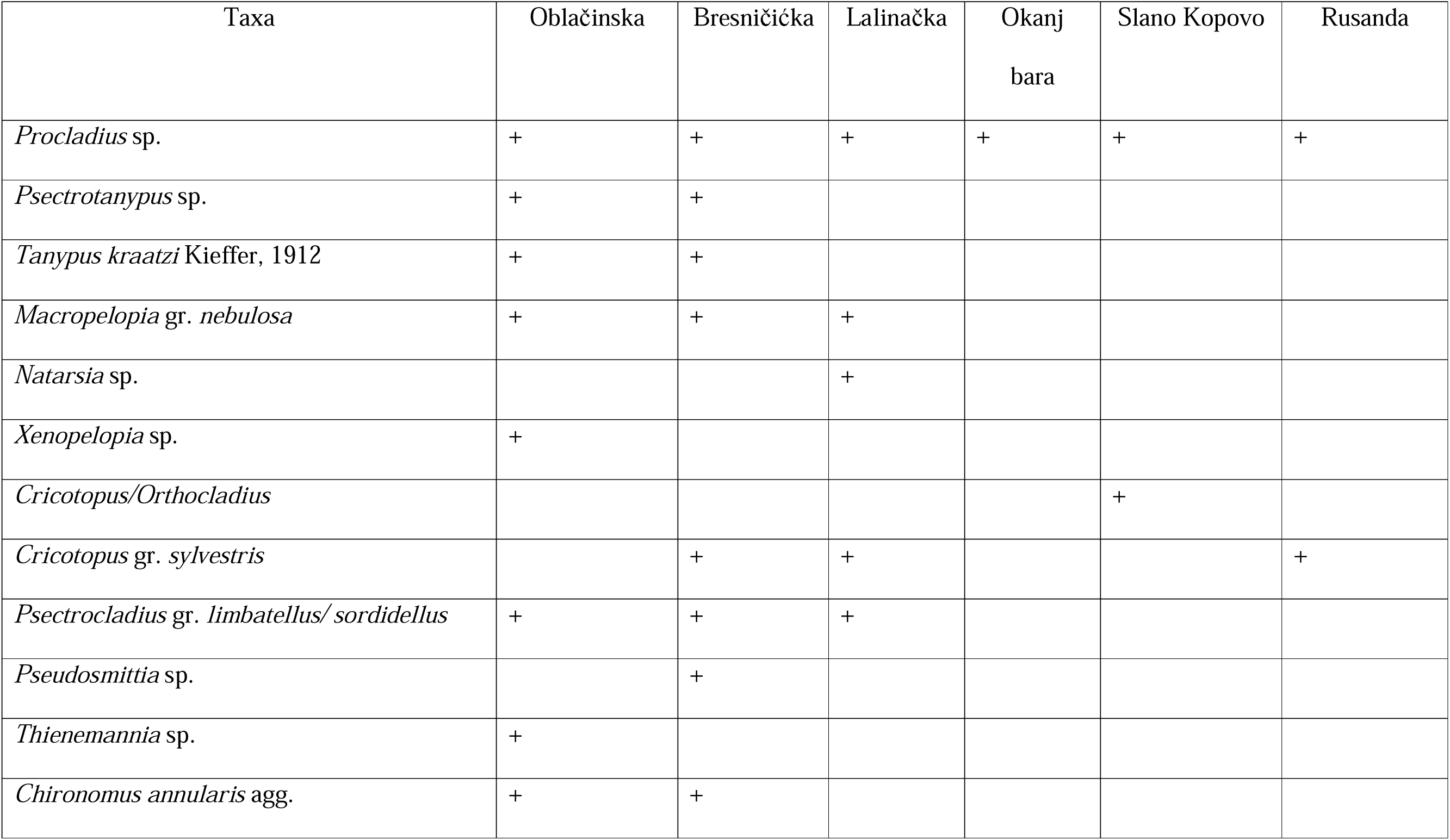
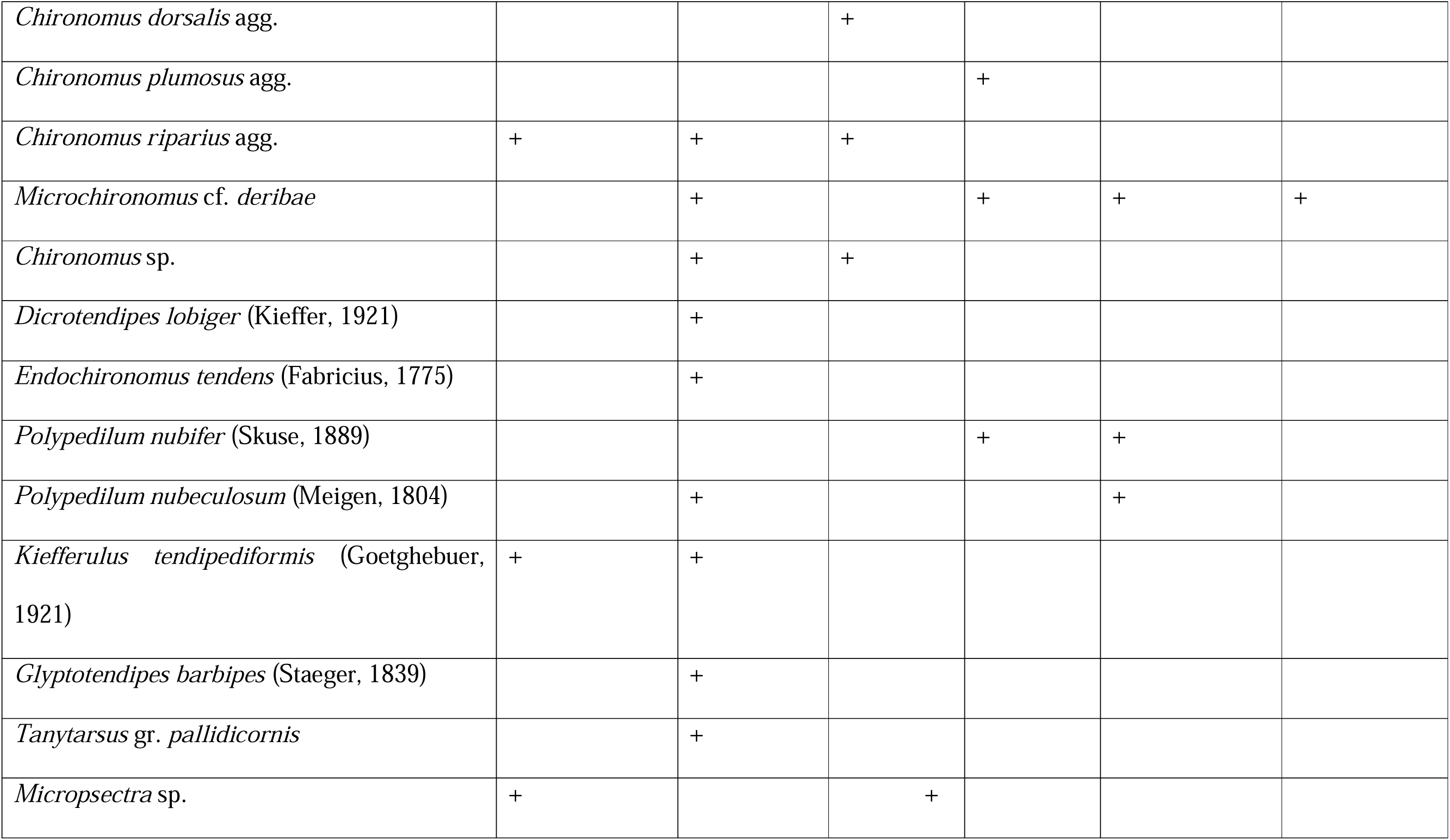
List of the recorded chironomid taxa across the study sites. + indicates the presence of the taxon.

### Seasonal dynamics of chironomid communities across the study sites

The highest species richness (S) and Shannon diversity (H) across all study sites were recorded in spring and summer months (Fig. 3), with exception of Oblačinska salt marsh where the highest species richness was recorded in winter (Fig. 3a). At Oblačinska salt marsh *Chironomus riparius* agg. dominated in abundance throughout the whole year (Fig. 4a, c and d) except during summer when *Chironomus annularis* agg. was the most abundant (Fig. 4b). *C. annularis* agg. was the most abundant at Bresničić salt marsh in summer and winter (Fig. 5b and d), *C. riparius* agg. dominated in abundance in spring (Fig. 5a), while *Microchironomus* cf. *deribae* dominated in autumn (Fig. 5c). *Procladius* sp. and *C. riparius* agg. were the most abundant taxa at Lalinačka salt marsh during spring (Fig. 6a). *C. dorsalis* agg. and *C. riparius* agg. dominated in abundance at Lalinačka salt marsh in autumn months (Fig. 6b), while *Natarsia* sp. was the only recorded taxon at this locality during winter months (Fig. 6c). As mentioned above, Lalinačka salt marsh was dry during whole summer in 2017. At both Okanj bara and Slano Kopovo chironomids were recorded only in summer. *Microchironomus* cf. *deribae* dominated in abundance at Okanj bara (Fig. 7a), while *Polypedium nubeculosum* was the most abundant taxon at Slano Kopovo (Fig. 7b). Finally, chironomids were recorded only in summer and autumn months at Rusanda site. *Procladius* sp. was the most abundant taxon during summer months (Fig. 7c), and it was the only recorded taxon in autumn (Fig. 7d).

**Fig. 3.**
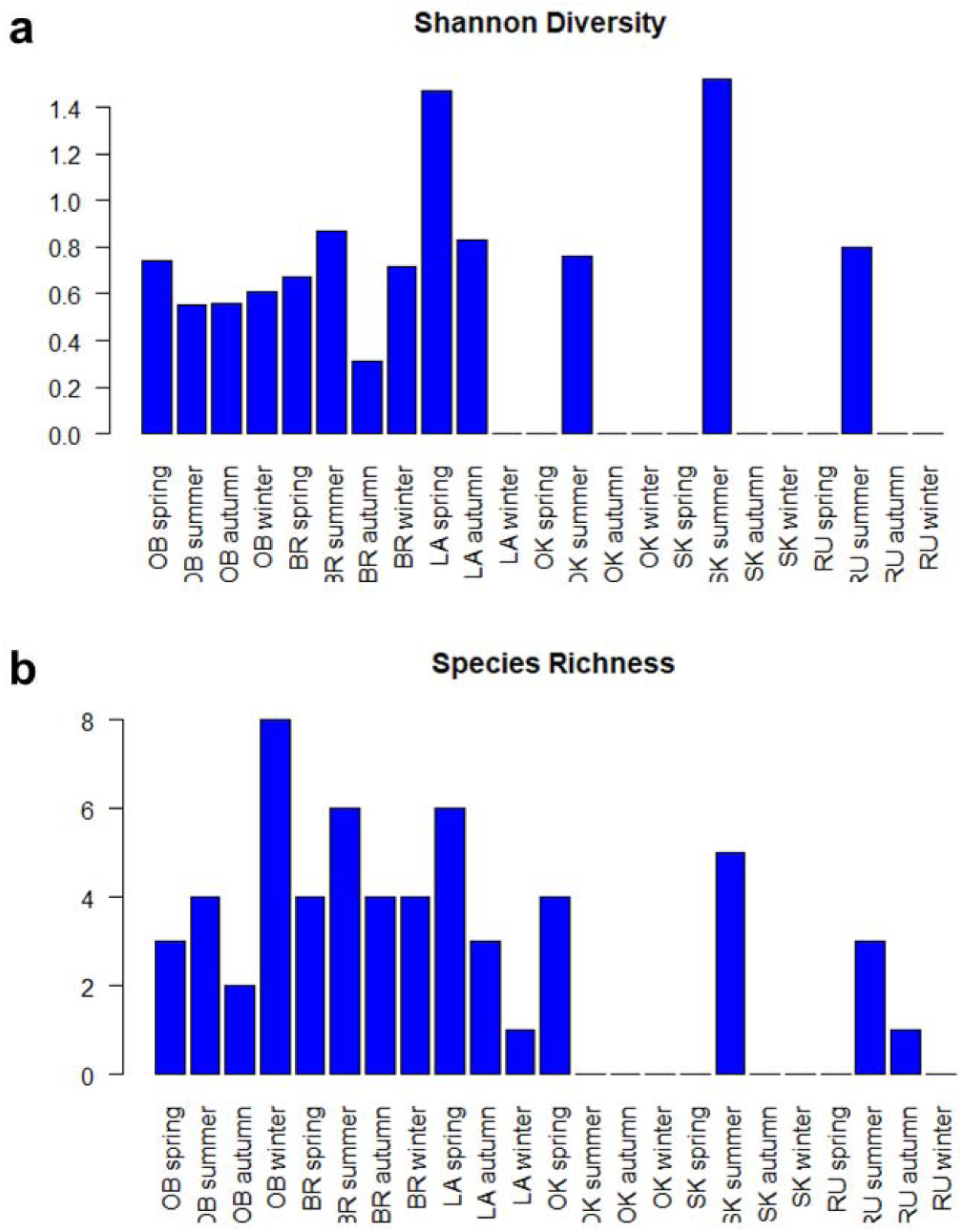
Variations of species richness (a) and Shannon diversity (b) across the study sites and through different seasons. Lalinačka salt marsh, LA; Oblačinska salt marsh, OB; Bresničićka salt marsh, BR; Okanj bara, OK; Slano Kopovo, SK; Rusanda, RU.

**Fig. 4.**
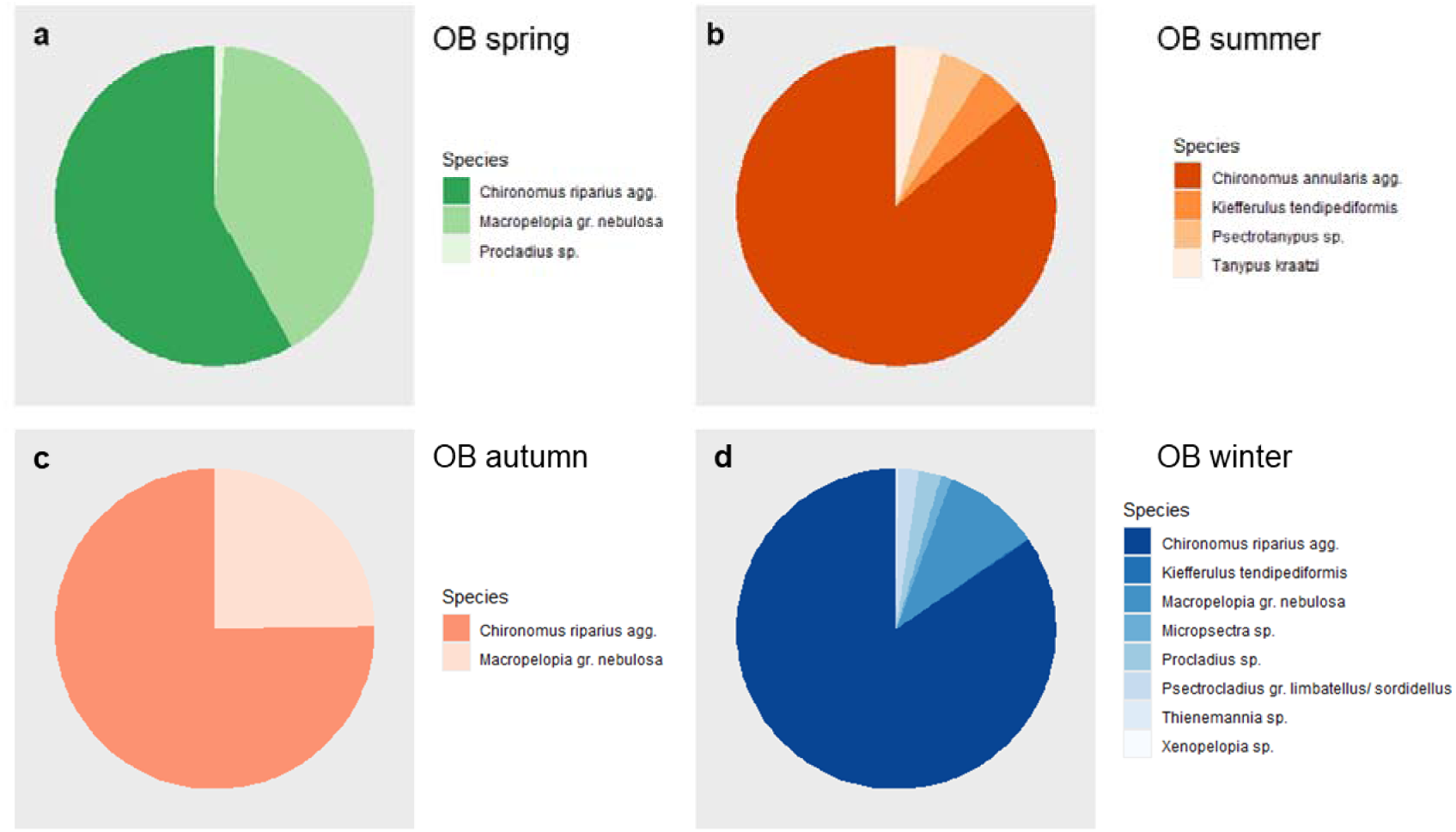
Community structure of chironomids at Oblačinska salt marsh (OB) in spring (a), summer (b), autumn (c), winter (d)

**Fig. 5.**
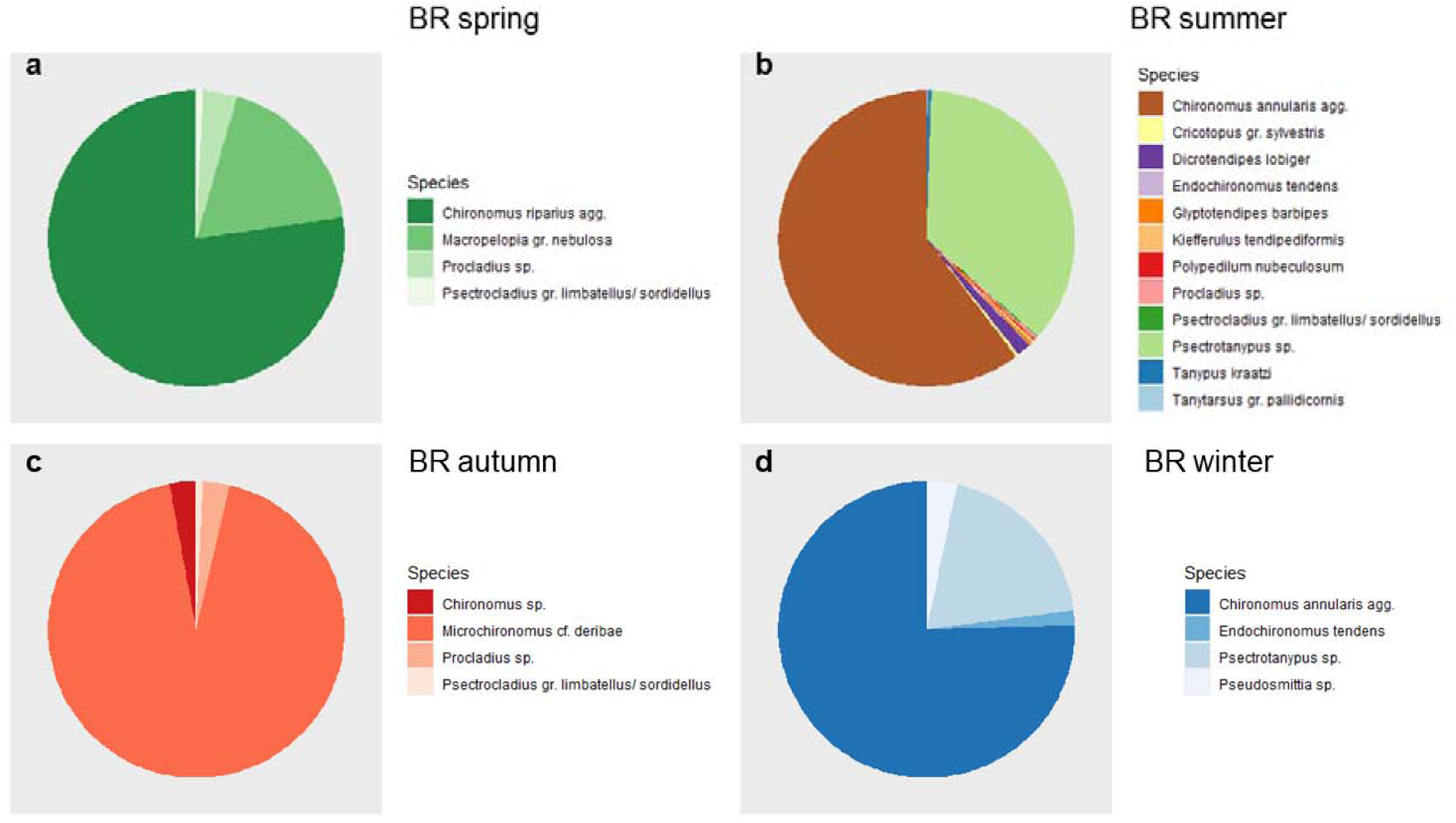
Community structure of chironomids at Bresničićka salt marsh (BR) in spring (a), summer (b), autumn (c), winter (d)

**Fig. 6.**
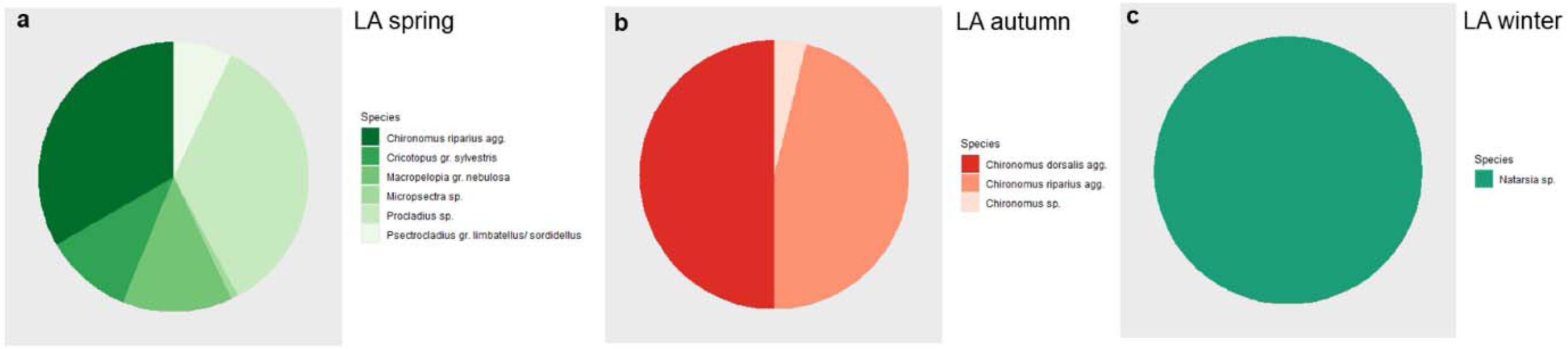
Community structure of chironomids at Lalinačka salt marsh (LA) in spring (a), autumn (b), winter (c)

**Fig. 7.**
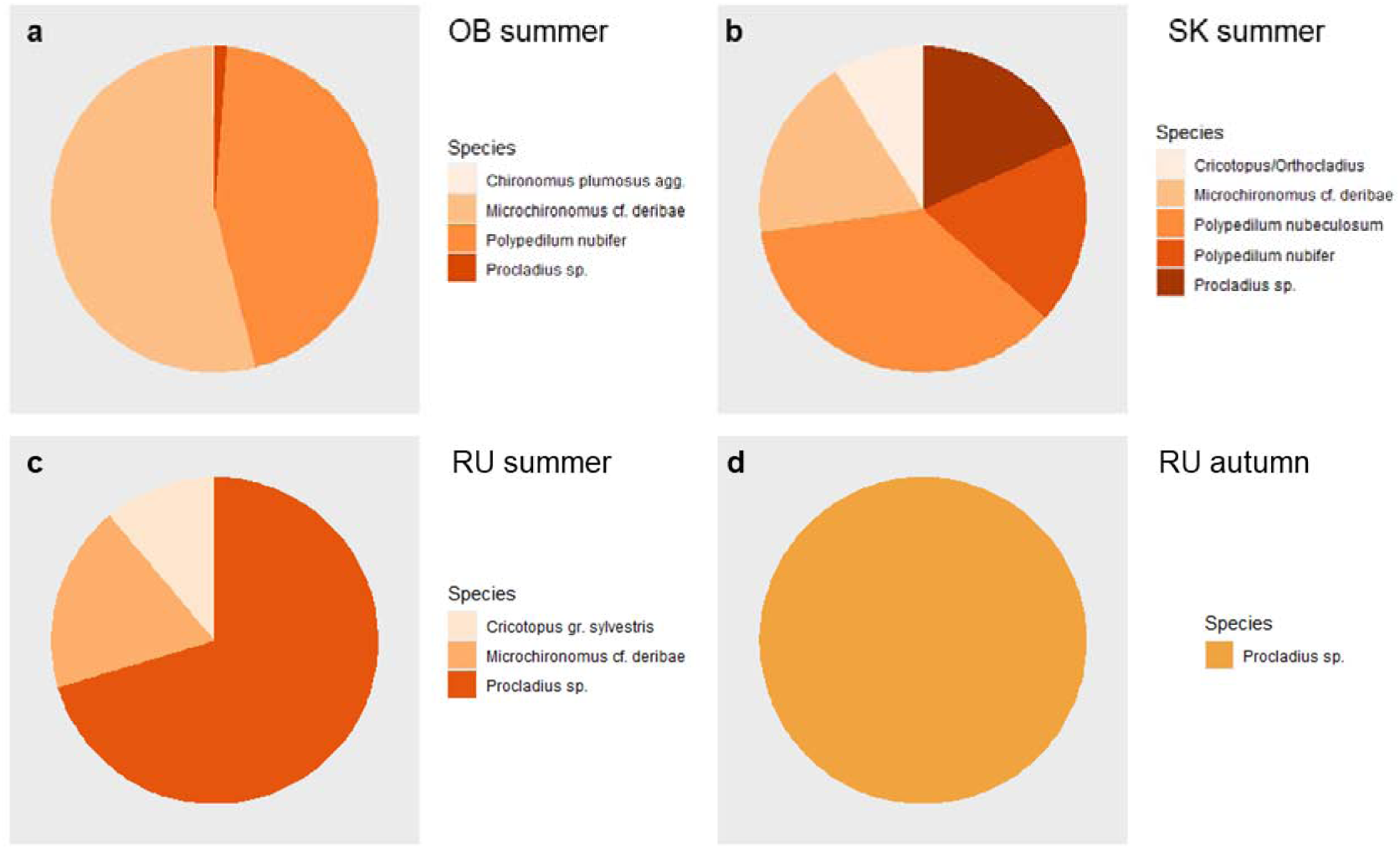
Community structure of chironomids at Okanj bara (OK) in summer (a), Slano Kopovo (SK) in summer (b), and Rusanda (RU) in summer (c) and autumn (d)

## Discussion

Our results indicate low species richness of chironomids in the Pannonian salt marshes in Serbia (from 3 to 5 taxa per study site) but higher species richness of chironomids in the salt marshes in south Serbia (from 9 to 17 taxa per study site). Also, we recorded a relatively low Shannon diversity (maximum 1.52) of the chironomid fauna at the salt marshes in Serbia. Similar results on chironomid species richness from benthic samples were recorded in freshwater ponds in a protected floodplain area in Croatia, where species richness varied from 4 to 19 taxa per study site (Čerba et al., 2023). The study of 22 urban waterbodies in Belgrade revealed a higher Shannon diversity of benthic chironomids in slow-flowing canals and reservoirs (mean Shannon diversity was around 2) while in urban rivers the Shannon diversity was around 3 (Popović et al., 2022). Lower diversity of chironomids in salt marshes from the northern part of the country comparing to the localities in the southern part of the country in our study is most probably the consequence of more extreme environmental conditions in Pannonian salt marshes. For instance, Lake Rusanda has the highest measured salinity of all saline lakes in the Carpathian Basin (Boros et al., 2014). Previous studies pointed out that species richness of aquatic macroinvertebrates tends to decrease with increased salinity in salt lakes due to the direct negative effects of high salinity or due to the decline in macrophyte diversity (Boda et al., 2019). In addition, the small surface of salt marshes in the southern part of the country in our study allows them to be completely overgrown by submerged macrophytes (Fig. 2a, b and c), which provides suitable conditions for higher number of chironomid species by increasing habitat complexity (Mormul et al., 2011).

Diversity (S and H) and community structure of chironomid communities in the studied salt marshes showed fluctuations related to the season. Seasonal variations in the structure of chironomid communities have often been observed (Milošević et al., 2013; Tóth et al., 2013; Głowacki et al., 2023), which is not surprising given that many environmental factors show cyclic variation during a year. The lowest diversity at the majority of study sites was recorded in winter. The exception is Oblačinska salt marsh where the highest species richness was recorded in winter. This locality dried up completely in late summer-early autumn, so the community started forming in the late autumn when the salt marsh was filled with water again. In the winter months, this locality reached the maximum water level. Earlier study of aquatic macroinvertebrates in temporary ponds showed that the diversity of macroinvertebrates was the highest during maximum water level and the lowest when ponds started filling with water (Bazzanti et al., 1996).

Most of the recorded chironomids in the studied salt marshes are common in freshwaters and are typical for eutrophic ecosystems. Interesting taxon recorded at Bresničićka salt marsh, *Pseudosmittia* sp. is semi-terrestrial or terrestrial (Moller Pillot, 2013). Some of the recorded taxa, such as *Chironomus annularis*, *C. plumosus*, *Microchironomus deribae*, *Polypedilum nubeculosum*, *Procladius* sp. and *Tanypus kraatzi* have been found to tolerate higher concentrations of chlorides in water (Moller Pillot & Vallenduuk, 2007; Moller Pillot, 2009). *Microchironomus* cf. *deribae* is the most tolerant to high salinity among all the recorded species (Moller Pillot, 2009) and, to our knowledge, this was the first record of this species for the chironomid fauna in Serbia, according to the published data (Janković, 1978, 1985; Milošević et al., 2011, 2018; Popović et al., 2016, 2022). This species has been recorded from many European countries along the seacoast (Moller Pillot, 2009), but also in soda pans in Hungary (Boda et al., 2019). The first record for chironomid fauna in Serbia here was also given for *Tanytarsus pallidicornis*. Other species from this genus have been recorded in Serbia, but in faunistic studies for chironomids in Serbia this taxon was identified only to the genus level (Milošević et al., 2011).

## Conclusions

Salt marshes in Serbia are poorly investigated, especially in terms of faunal composition. Our study contributes to the knowledge of their biota and to the knowledge of the chironomid fauna in Serbia which are important components of aquatic habitats, including salt marshes. Overall, we recorded a total of 25 taxa of which two are new for the Serbian chironomid fauna. The present findings set the basis for future research and conservation measures in these unique and fragile habitats.

## Acknowledgements

This study was supported by grant 18197-1: Establishing conservation management of salt marshes in Serbia based on monitoring of macroinvertebrate community, funded by the Rufford Foundation, UK, and by Grant No. 451-03-66/2024-03/200124, funded by the Serbian Ministry of Science, Technological Development, and Innovation, for O.S. We thank students from the Department of Biology and Ecology, Faculty of Sciences and Mathematics Niš, and members of the Biological Society “Dr. Sava Petrović” from Niš for helping in the field and laboratory work. We would also like to thank Dr. Ana Savić and Dr. Djuradj Milošević from the Faculty of Sciences and Mathematics, University of Niš, for their help in organizing the study and early stages of the field survey.

## Notes

### Competing Interest Statement

The authors have declared no competing interest.

